# Surveying the vampire bat (*Desmodus rotundus*) serum proteome: a resource for identifying immunological proteins and detecting pathogens

**DOI:** 10.1101/2020.12.04.411660

**Authors:** Benjamin A. Neely, Michael G. Janech, M. Brock Fenton, Nancy B. Simmons, Alison M. Bland, Daniel J. Becker

## Abstract

Bats are increasingly studied as model systems for longevity and as natural hosts for some virulent viruses. Yet our ability to characterize immune mechanisms of viral tolerance and to quantify infection dynamics in wild bats is often limited by small sample volumes and few species-specific reagents. Here, we demonstrate how proteomics can overcome these limitations by using data-independent acquisition-based shotgun proteomics to survey the serum proteome of 17 vampire bats (*Desmodus rotundus*) from Belize. Using just 2 μL of sample and relatively short separations of undepleted serum digests, we identified 361 proteins across five orders of magnitude. Data are available via ProteomeXchange with identifier PXD022885. Levels of immunological proteins in vampire bat serum were then compared to human plasma via published databases. Of particular interest were anti-viral and anti-bacterial components, circulating 20S proteasome complex, and proteins involved in redox activity; whether any results are specific to vampire bats could be assessed by future pan-mammalian analyses. Lastly, we used known virus proteomes to identify Rh186 from *Macacine herpesvirus 3* and ORF1a from Middle East respiratory syndrome-related coronavirus, indicating that mass spectrometry-based techniques show promise for pathogen detection. Overall, these results can be used to design targeted mass-spectrometry assays to quantify immunological markers and detect pathogens. More broadly, our findings also highlight the application of proteomics in advancing wildlife immunology and pathogen surveillance.

## Introduction

Bats are a hyper-diverse and geographically widespread order (Chiroptera), accounting for over 20 % of all mammal species^1^. Owing to their capacity for flight and wide range of trophic habits (*e.g*., frugivory, nectarivory, insectivory, carnivory, sanguivory), bats provide critical ecosystem services that include seed dispersal, pollination, and predation of insects ^2^. Unique features of bats among mammals, such as the capacity for powered flight and long lifespans despite small body sizes, also make this order interesting for basic research in ecology and evolution ^3,4^. Recently, bats have also become model systems for studies of the microbiome and sociality^5,6^.

However, because some bats are common in anthropogenic landscapes, they are increasingly studied for their ability to harbor pathogens with high virulence in humans and domestic animals ^7,8^. In particular, bats carry more zoonotic viruses than most other mammalian orders ^9^, and they are the confirmed reservoir hosts for Hendra virus, Nipah virus, Marburg virus, an array of lyssaviruses (*e.g*., rabies virus), and SARS-like coronaviruses ^10–13^. Spillovers of these sometimes lethal viruses, from bats to humans, are often driven by ecological changes that alter infectious disease dynamics in wild bat populations and change interactions between bats and recipient hosts ^14,15^. However, with some exceptions (*e.g*., lyssaviruses), these viruses may not typically cause clinical disease in bats ^16^. Although the high richness of zoonotic viruses in bats may simply be a function of their vast species diversity ^17^, tolerance to virulent viruses is likely driven by distinct aspects of immunity in these flying mammals, including robust complement, constitutive expression of type I interferons (*e.g*., IFN-α), and high combinatorial diversity in immunoglobulin genes ^4,18–20^. Accordingly, contemporary studies of zoonotic pathogens in bats have focused on characterizing immunological factors that facilitate tolerance or shedding of viruses, quantifying viral diversity, and identifying spatial and temporal pulses of infection ^21–23^.

Traditionally, efforts to assess the immunological state of wild bats and detect viruses have relied on relatively simple tools. Most bat species are sufficiently small that only modest volumes of blood can be safely collected under non-lethal procedures ^24^; 74% of bat species weigh less than 30 grams as adults ^25^. Small blood volumes and lack of species-specific reagents for bats generally restrict the scope of possible immune assays ^26,27^. Remote field sites also pose significant challenges for sample storage and transport at ideal temperatures. Despite these obstacles, techniques such as *in vitro* microbial killing assays, enzyme-linked immunosorbent assays, and *in vivo* antigen challenge have helped to broadly characterize complement, antibody, and cellular immune response in wild bats and how these phenotypes vary with life history (*e.g*., reproduction) and environmental conditions ^28–30^. Similarly, polymerase chain reaction (PCR) has formed the primary basis of virus detection ^31^. However, because bat colonies can be large (e.g., hundreds to over 20 million individuals ^32^) and prevalence of active viral infection is generally low, serology and the detection of virus-specific antibodies has also commonly been employed ^33^. Although serological surveys have been important for characterizing virus circulation in bats, antibody cross-reactivity and inconsistent cut-off thresholds can limit their interpretability ^34^.

Given these restrictions, studies of bat immunology and virology have increasingly benefitted from modern bioanalytical approaches. Researchers employing global profiling techniques, such as RNA-Seq-based transcriptomics, have begun to contribute new resources for bat immunology and to illuminate the broad immune response of bats to viral infections ^35,36^. Similarly, metagenomic approaches have helped reduce bias in broad characterization of bat viral diversity ^37,38^. In addition to these approaches, proteomics can provide a complementary modality to define the molecular landscape. Proteomics is uniquely useful when applied to blood, because the blood proteome is mostly secreted from organs such as the liver ^39^, thereby permitting interrogation of the circulating protein phenotype beyond cellular transcript profiling. Using the relatively small sample volumes that typify bat field studies (*e.g*., < 10 μL), proteomic analysis of blood can identify and relatively-quantify hundreds of proteins, including but not limited to those involved in host response to infection and other useful immunological biomarkers ^40^. Prior applications of serum proteomics in bats have described proteins involved in shifts in innate immunity and blood coagulation between hibernating and active greater mouse-eared bats (*Myotis myotis*)^41^ as well as between North American and European *Myotis* species that vary in infection with *Pseudogymnoascus destructans*, which causes white-nose syndrome ^42^. Proteomic applications to other bat tissues, such as saliva, brain, and lung, have identified possible immunomodulatory properties in vampire bats (*Desmodus rotundus*) ^43^, neuronal differences between active and torpid horseshoe bats (*Rhinolophus ferrumequinum*) ^44^, and upregulation of cell-mediated immunity in flying foxes (*Pteropus alecto*) compared with ferrets infected with Hendra virus ^45^. Given the general utility of proteomic analysis, we expected a survey of the vampire bat serum proteome should provide complementary information to existing approaches, while also providing a more complete survey of the underlying molecular landscape.

Here, we used data-independent acquisition-based shotgun proteomics (i.e., bottom-up proteomics) to profile the undepleted serum proteome of 17 bats from two locations. We focused this work on vampire bats, a species that has an obligate diet of blood and feeds on prey as diverse as sea lions, tapirs, livestock, and humans ^46–48^, providing vast opportunities for transmission of viruses (e.g., rabies virus, adenovirus, herpesvirus) to and from these prey ^49–51^. Our results demonstrate the feasibility and capabilities of serum proteomic analyses in wild bats, including possibilities to simultaneously detect immunological components and viruses as well as to establish preliminary ranges of vampire bat proteins for comparison with other mammals.

## Methods

### Vampire bat sampling

As part of an ongoing longitudinal study, vampire bats were sampled in 2015 at two adjacent localities in the Orange Walk District of Belize: Lamanai Archeological Reserve (LAR, 450 hectares) and Ka’Kabish (KK, 45 hectares). These sites are located in a highly agricultural mosaic landscape where deforestation has been driven by cropland expansion in the 1990s followed by heightened demand for livestock pasture in the 2000s ^52^. These land-use changes have fragmented previously intact forest and have provided vampire bats with an abundant prey in the form of livestock ^29^. In recent years (2016 onward), rabies outbreaks in domestic animals have increased across Belize and in Orange Walk District, and virus isolates from livestock have been characterized as vampire bat-associated variants ^53^. Although we are unaware of other viral detection efforts in vampire bats in Belize, this species has tested positive elsewhere in Central and South America for adenoviruses, coronaviruses, flaviviruses, hantaviruses, herpesviruses, and paramyxoviruses, as well as for antibodies against henipavirus-like viruses ^51,54–62^.

Vampire bats were captured with mist nets and harp traps set along trails and adjacent to known roost sites. All individuals were issued a unique incoloy wing band (3.5 mm, Porzana Inc.) and identified by sex, age, and reproductive status ^63^. Serum was obtained by lancing the propatagial vein with a 23-gauge needle to collect blood and centrifuging resulting samples in serum separator tubes followed by short-term freezing at −20 °C until long-term −80 °C storage. Bleeding was stopped with styptic gel, and all bats were released at their capture location. Field protocols followed guidelines for safe and humane handling of bats issued by the American Society of Mammalogists ^24^ and were approved by the University of Georgia Animal Care and Use Committee (A2014 04-016-Y3-A5). Bat sampling was authorized by the Belize Forest Department under permit number CD/60/3/15(21). Specimen use for proteomic analysis was approved by the NIST Animal Care and Use Coordinator (NIST ACUC) under approval MML-AR19-0018. We included 17 serum samples for proteomic analysis, from 11 bats sampled at LAR (10 males, one female) and six bats sampled at KK (five males, one female). This information and the experimental key can be found in Supplemental Table S1.

### Sample processing and digestion

Prior to analysis, the sample set was randomized and assigned an experimental key to avoid bias and minimize batch effects. The 17 serum samples were thawed then centrifuged at 13 000 x *g*n for 8 min at 4 °C. The S-Trap method was used for digestion with the S-Trap micro column (ProtiFi; ≤ 100 μg binding capacity), and the specific method is described below. Based on prior work with other mammalian sera, we assumed a protein concentration of approximately 50 μg/μL (this value can vary but assuming at least 50 μg/μLwas used for estimating enzyme for the digest). Therefore, 2 μL (approximately 100 μg protein) of each sample was mixed with 48 μL 50 mmol/L ammonium bicarbonate and 50 μL of 2x lysis buffer (ProtiFi), then added onto the STrap. Additionally, since the samples were digested in two sets, each set included a 2 μL aliquot of NIST SRM 909c Frozen Human Serum, which is technically a converted plasma pool. This sample was used to qualitatively confirm digestion, LC-MS/MS performance, and downstream search methods. All samples were reduced with 10 μL of 90 mmol/L DL-Dithiothreitol (DTT; final concentration of 10 mmol/L) at 60 °C for 30 min, then cooled and alkylated with 10 μL of 200 mmol/L 2-chloroacetamide (CAA; final concentration of 20 mmol/L) at room temperature in the dark for 30 min. The sample was acidified with 12 μL of 12 % phosphoric acid (volume fraction) bringing the final volumetric ratio to 1:10. Next, 700 μL binding buffer [ProtiFi; 5 % triethylammonium bicarbonate (TEAB) volume fraction in methanol] was added (approximately 1:7 volumetric ratio). Using a vacuum manifold, the sample was washed across the S-Trap with six sequential washes of 400 μL binding buffer. Next, 3 μL of 1 μg/μL trypsin (Pierce) was mixed with 122 μL, 50 mmol/L ammonium bicarbonate, and this 125 μL solution was added to each S-Trap, yielding approximately a 1:30 mass ratio (trypsin:total protein). Samples were incubated at 47 °C for 1 h, after which they were sequentially washed into 1.5 mL Lo-Bind microcentrifuge tubes (Eppendorf) by centrifugation with the following wash steps: 80 μL 50 mmol/L ammonium bicarbonate, 80 μL 0.2 % formic acid (volume fraction), 80 μL 0.2 % formic acid in 50 % acetonitrile (volume fractions) at 1000 x *g*n, 1000 x *g*n, and 4000 x *g*n, respectively at 4 °C for 1 min. The resulting peptide mixtures were reduced to dryness in a vacuum centrifuge at low heat and stored at −80 °C. Prior to analysis samples were reconstituted with 100 μL 0.1 % formic acid (volume fraction) and briefly vortexed, then centrifuged 10 000 x *g*n for 10 min at 4 °C. The peptide concentration of each sample was determined using the Pierce quantitative fluorometric peptide assay with a BioTek Synergy HT plate reader.

### LC-MS/MS

Peptide mixtures in 0.1% formic acid (volume fraction) were analyzed using an UltiMate 3000 Nano LC coupled to a Fusion Lumos Orbitrap mass spectrometer (Thermo Fisher Scientific). Using the original sample randomization yielded a randomized sample order and injection volumes were determined for 0.5 μg loading (between 0.37 and 0.95 μL sample). One sample from bat D141 (experimental key Bat_20) was used as a technical replicate to evaluate technical variability across the run (Supplemental Figure S1), while the two aforementioned human serum pools were analyzed in the same manner as the bat sera. Peptide mixtures were loaded onto a PepMap 100 C18 trap column (75 μm id x 2 cm length; Thermo Fisher Scientific) at 3 μL/min for 10 min with 2 % acetonitrile (volume fraction) and 0.05 % trifluoroacetic acid (volume fraction) followed by separation on an Acclaim PepMap RSLC 2 μm C18 column (75μm id x 25 cm length; Thermo Fisher Scientific) at 40 °C. Peptides were separated along a 60 min two-step gradient of 5 % to 30 % mobile phase B (80 % acetonitrile volume fraction, 0.08 % formic acid volume fraction) over 50 min followed by a ramp to 45 % mobile phase B over 10 min and lastly ramped to 95% mobile phase B over 5 min, and held at 95 % mobile phase B for 5 min, all at a flow rate of 300 nL/min.

The Fusion Lumos was operated in positive polarity with a data-independent acquisition (DIA) method constructed using the targetedMS2 module (as opposed to the built-in DIA module). The full scan resolution using the orbitrap was set at 120 000, the mass range was 399 to 1200 *m/z* (corresponding to the DIA windows used), 30 % RF lens was set, the full scan ion target value was 4.0e5 allowing a maximum injection time of 20 ms. A default charge of 4 was set under MS Global Settings. As stated, DIA windows were constructed using the targetedMS2 module. Each window used higher-energy collisional dissociation (HCD) at a normalized collision energy of 32 with quadrupole isolation width at 21 *m/z*. The fragment scan resolution using the orbitrap was set at 30 000, the scan range was specified as 200 to 2000 *m/z*, ion target value of 1.0e6 and 60 ms maximum injection time. Data were collected as profile data in both MS1 and MS2, though the authors wish to note that using centroid data is possible for most DIA software. The DIA window scheme was an overlapping static window strategy such that each of the 40 windows were 21 *m/z* wide, with 1 *m/z* overlap on each side covering the range of 399 to 1200 *m/z*. The window centers were specified in the mass list table (with z = 2), such that they were 409.5, 429.5, 449.5,…, 1189.5. The method file (85min_DIA_40×21mz.meth) is also included in the PRIDE submission. The mass spectrometry proteomics data have been deposited to the ProteomeXchange Consortium via the PRIDE ^64^ partner repository with the dataset identifier PXD022885.

### DIA search parameters

Analysis was performed using Spectronaut (v13.6.190905.43655). The following settings were used. Sequences: Trypsin selected, max pep length 52, min pep length 7, two missed cleavages, KR as special amino acids in decoy generation, toggle N-terminal M set to true. Labelling: no labelling settings were used. Applied modifications: maximum of five variable modifications using fixed carbamidomethyl (C), and variable acetyl (protein N-term) and oxidation (M). Identification: per run machine learning, Q-value cut-off of 0.01 for precursors and proteins, single hits defined by stripped sequence, and do not exclude single hit proteins, PTM localization set to true with a probability cut-off of 0.75, kernel density p-value estimator. Quantification: interference correction was used with excluding all multi-channel interferences with minimum of 2 and 3 for MS1 and MS2, respectively, proteotypicity filter set to none, major protein grouping by protein group ID, minor peptide grouping by stripped sequence, major group quantity set to mean peptide quantity, a Major Group Top N was used (min of 1, max of 3), minor group quantity set to mean precursor quantity, a Minor Group Top N was used (min1, max of 3), quantity MS-level used MS2 area, data filtering by q-value, cross run normalization was used with global median normalization and automatic row selection, no modifications or amino acids were specified, best N fragments per peptide was set to between 3 and 6, with ion charge and type not used. Workflow: no workflow was used. Post Analysis: no calculated explained TIC or sample correlation matrix, differential abundance grouping using major group (from quantification settings) and smallest quantitative unit defined by precursor ion (from quantification settings), differential abundance was not used for conclusions. The fasta file used for searching bat samples was the NCBI RefSeq *Desmodus rotundus* Release 100, GCF_002940915.1_ASM294091v2 (29 845 sequences), and for searching human samples was the 2020_01 release of the UniProtKB SwissProt with isoforms, using the taxonomy term 9606 (*Homo sapiens*; 42 385 sequences). For the host plus virus searching, these same fasta files were used along with a fasta of *D. rotundus*–associated virus sequences retrieved from the DBatVir database (156 sequences; downloaded 12 November 2020 ^62^) and the following six fasta retrieved from the 2020_05 release of the UniProtKB SwissProt and TrEMBL: Adenoviridae (taxon ID: 10508; 30 051 sequences), *Betacoronavirus* (taxon ID: 694002; 21 892 sequences), *Henipavirus* (taxon ID: 260964; 462 sequences), Herpesvirales (taxon ID: 548681; 88 353 sequences), *Morbillivirus* (taxon ID: 11229; 23 142 sequences), and *Rabies lyssavirus* (taxon ID: 11292; 23 398 sequences). All fasta and Spectronaut .snes files (the human and bat data are searched with and without all virus databases) are included in PRIDE submission PXD022885.

### Ortholog mapping, gene ontology term sorting, and rank comparisons

Following DIA identification, there were 376 RefSeq vampire bat protein identifiers with MS2-based quantities across the experiment (Supplemental Table S2). These identifications were converted to human orthologs to aid in downstream analysis and for comparison to other studies. This was accomplished by using a series of python scripts (Anaconda v2019.07; conda v4.7.11; Python v3.6.8) from Github on the following two repositories: pwilmart/PAW_BLAST and pwilmart/annotations, retrieved November 18, 2019. Broadly, these tools take a list of identifiers to create a subset fasta (make_subset_DB_from_list_3.py), which is then searched against a human fasta (db_to_db_blaster.py) using a local installation of BLAST+ 2.9.0 ^65^. The results were further annotated using add_uniprot_annotations.py. The 376 vampire bat proteins were assigned human orthologs, which were then manually inspected for incorrect assignments. Such assignments can happen when a protein is not present in humans (*e.g*., beta-lactoglobulin 1 or inhibitor of carbonic anhydrase), or if the blast hit disagrees with the original annotation, in which case the original annotation is preserved (*e.g*., apolipoprotein R was matched to C4b-binding protein alpha chain). Finally, duplicate entries were summed together into a single non-redundant entry. An intermediate table (Supplemental Table S2) is given that delineates which entries had manually assigned gene names, or were duplicates to be summed. The resulting table included mapped orthologs with UniProtKB links (Supplemental Table S3). Gene ontology (GO) term sorting was performed by using GO search terms in UniProtKB along with the human taxon identifier (9606), yielding a list of human gene symbols that was used to subset the vampire bat results. Specifically in Supplemental Table S3, the complement activation Gene Ontology (GO) term (GO:0006956) was used to identify 31 bat proteins. In addition to vampire bat proteome, identifications and relative abundances of the control pooled human serum can be found in Supplemental Table S4, but were not used for comparison.

The serum proteome of the vampire bat was compared to the plasma proteome of the human described on the Human Blood Atlas by using values from Table 1 “Protein detected in human plasma by mass spectrometry” from the Human Blood Atlas website (https://www.proteinatlas.org/humanproteome/blood/proteins+detected+in+ms; accessed 15 June 2020). This table was combined with the relative abundance and resulting ranks of the 361 proteins identified (Supplemental Table S5). The rank of the shared 323 proteins was compared by determining the absolute difference in rank divided by the minimum rank, referred to as “Rank Delta”. An arbitrary cut-off of 3 was chosen, but the complete list was still manually inspected for proteins of interest. Only proteins that were exceptionally elevated (or closely related to those that were elevated) in the vampire bat serum proteome were considered. Some proteins like PRDX1 (136th in bat and 432 in human) were less than 3 in their rank delta, but were still considered of interest. This final list of proteins of interest can be found in Supplemental Table S6, along with links to both UniProt and Human Blood Atlas entries.

## Results and Discussion

Studying the interactions between hosts and pathogens is important to understand infectious disease risks, yet such research is hampered by our poor knowledge of the underlying biochemical landscape for most host species. Bats offer a unique opportunity to investigate how immunological components may confer the ability to harbor virulent viruses, a topic of considerable interest particularly in light of the current COVID-19 pandemic. Because complement plays a key role in innate immunity and functions as a bridge between innate and adaptive responses ^66–68^, prior studies have mostly used microbicidal assays using pathogens with complement-dependent killing to profile levels of these proteins in bats, demonstrating individual and environmental sources of variation ^28–30,69,70^. We profiled the serum proteome of vampire bats as a first step at broadly describing the bat serum proteome. This inventory allowed investigating not only complement proteins, but also a global investigation of the bat serum proteome, including other components of the innate immune response, as well as detection of viral proteins, and can serve as a resource to develop targeted mass spectrometry-based assays.

Blood is a unique biofluid that is proximal to most organs, containing cells and secreted proteins. The considerable dynamic range of proteins in serum and plasma has been frequently described in humans, spanning at least eight orders of magnitude, though approximately 10 proteins account for 90 % of the plasma proteome ^39,71^. Similarly, we unsurprisingly found the vampire bat serum proteome encompassed a dynamic range of five orders of magnitude (Figure 1A). The top four proteins (albumin, immunoglobulin lambda-1 light chain, hemoglobin subunit beta, and alpha-1-antitrypsin) accounted for 50% of the total abundance estimated using DIA. Ranges for the top 20 proteins (based on average abundance; Figure 1B) were determined using protein abundance from the 17 individuals. Despite the difficulties in making direct comparisons between the vampire bat serum proteome and that of other mammals, it is possible to specifically evaluate complement components in bat serum. The complement activation Gene Ontology (GO) term (GO:0006956), which includes the lectin pathway, was used to highlight relative levels of 31 proteins in bat serum (Figure 1C) and indicate their rank and abundance in the serum proteome (Figure 1A). Apolipoprotein R and manose-binding protein A were added since they are not present in humans but are likely involved in complement activation. It would be interesting to infer that these complement protein levels are higher in bats than other mammals, but without adequate reference ranges, and given the technical artifacts of serum preparation across species, these comparisons are currently difficult at best and an avenue for future comparative study.

**Figure 1.**
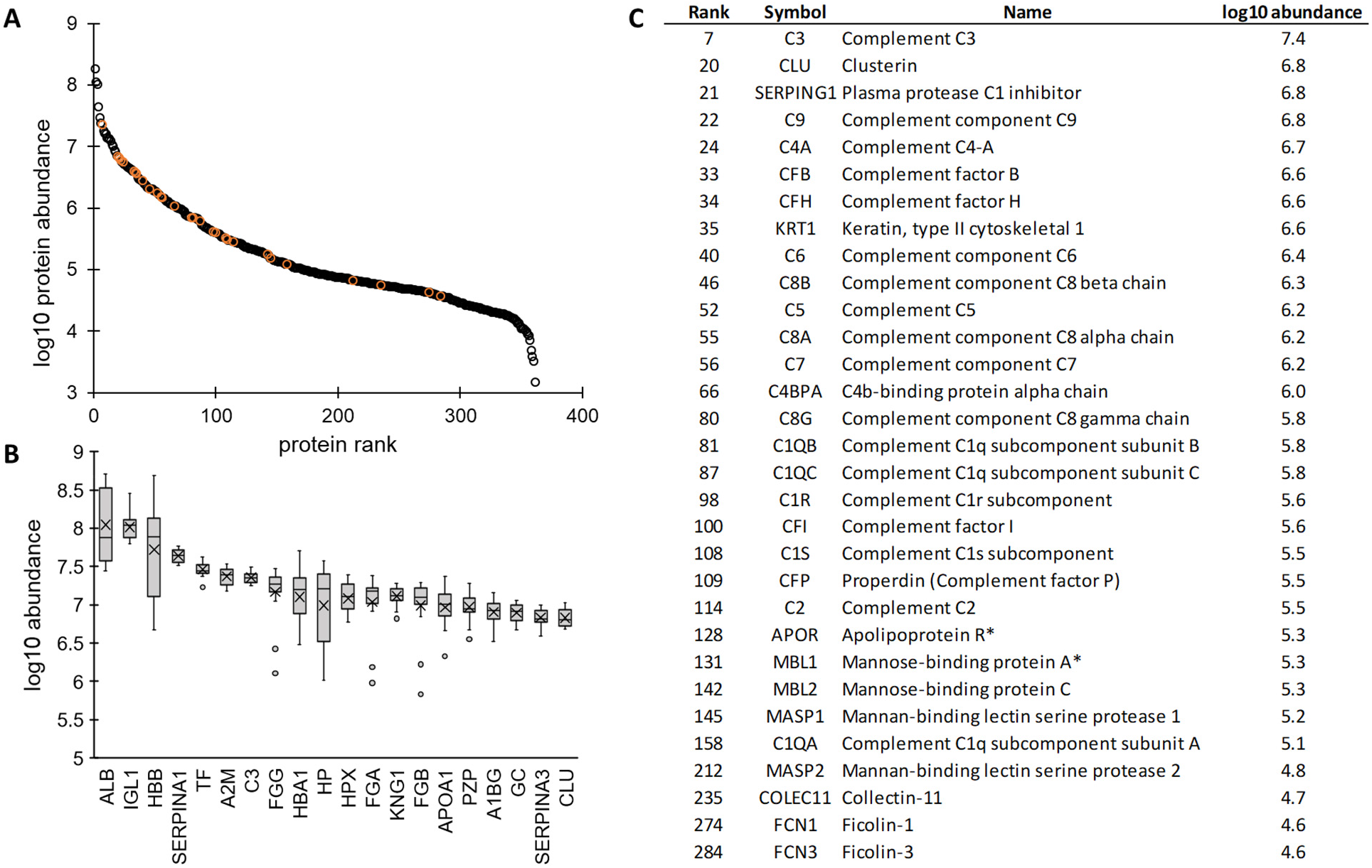
Average serum protein abundance. **A.** The average intensity of the 361 identified proteins was plotted with rank to illustrate the dynamic range of the vampire bat serum proteome. Complement related proteins are in orange for reference. **B.** Box and whisker plot of the top 20 proteins, listed as gene symbols: ALB, serum albumin; IGL1, immunoglobulin lambda-1 light chain; HBB, hemoglobin subunit beta; SERPINA1, alpha-1-antitrypsin; TF, serotransferrin; A2M, alpha-2-macroglobulin; C3, complement C3; FGG, fibrinogen gamma chain; HBA1, hemoglobin subunit alpha; HP, haptoglobin; HPX, hemopexin; FGA, fibrinogen alpha chain; KNG1, kininogen-1; FGB, fibrinogen beta chain; APOA1, apolipoprotein A-I; PZP, pregnancy zone protein; A1BG, alpha-1B-glycoprotein; GC, vitamin D-binding protein; SERPINA3, alpha-1-antichymotrypsin. **C.** 31 proteins involved in complement activation (GO:0006956). The rank and abundance are given for easy reference to the A panel. The ‘*’ for MBL1 and APOR is to indicate that these are not human proteins but likely involved in complement activation.

Although blood has been extensively studied in humans, it is surprisingly difficult to compare mass spectrometry-based analysis between human studies, much less between different species. The current study used a pooled human converted plasma sample (Supplemental Table S4) to enable this comparison. Additionally, the Human Blood Atlas ^72^ was used as a reference (Supplemental Table 5), with reported approximate concentrations of over 3000 proteins detected by mass spectrometry in human plasma. For an exploratory analysis, using these two datasets is acceptable, but two key caveats should be acknowledged. First, ranges of human concentrations are needed, which is obscured in a pooled sample as well as in the current Human Blood Atlas. Second, the wild bat samples may have had technical artifacts (e.g., hemolysis), which is evident by the high levels of hemoglobin subunit beta likely resulting from ruptured red blood cells ^73^ (Figure 1B). Ongoing efforts by our group and others will continue to define the serum and plasma proteome in humans and other species, and we expect more population-level data with protein abundance ranges to become available and more easily comparable across mammals.

Despite these caveats, we can highlight certain proteins that are higher in bat serum than in humans, without an obvious technical artifactual explanation. These were identified using the ranks of proteins in the parallel analyzed human pool and those available on the Human Blood Atlas versus the ranks of proteins in the vampire bat dataset (Figure 2; Supplemental Table S5). Broadly, these proteins fall into four general categories: innate immunity, circulating proteasome, anti-oxidants, and “other”. We encourage readers to explore the full table for specific comparative proteins of interest (Supplemental Table S6).

**Figure 2.**
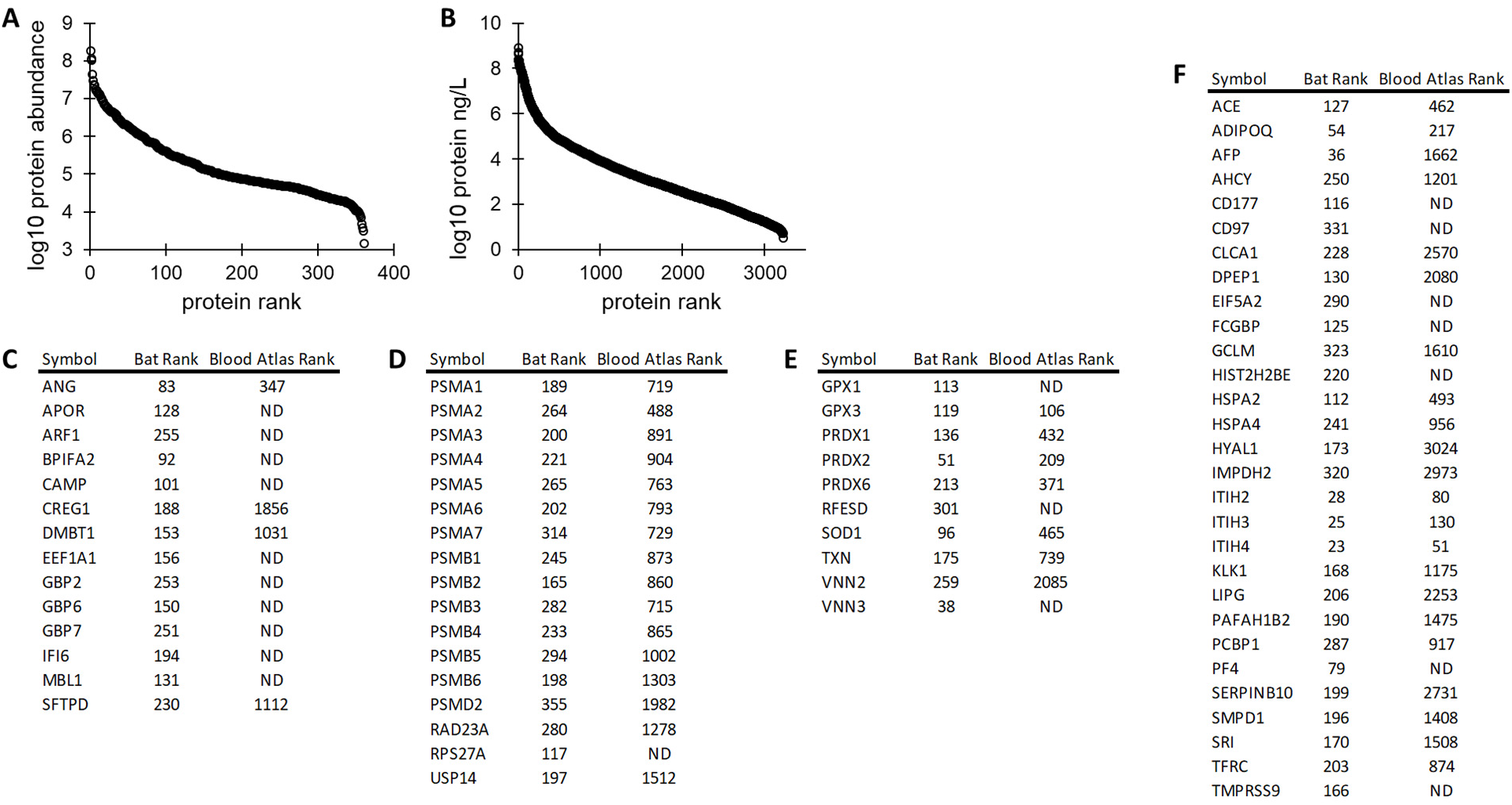
Proteins elevated in vampire bat serum versus human plasma. **A.** Average protein abundance of 361 proteins in vampire bat serum for reference. **B.** Reported protein abundance of 3222 proteins reported on the Human Plasma Atlas, for reference. **C.** Innate immunity proteins with bat and human ranks. ANG, angiogenin; APOR, apolipoprotein R; ARF1, ADP-ribosylation factor 1; BPIFA2, BPI fold-containing family A member 2; CAMP, cathelicidin antimicrobial peptide; CREG1, protein CREG1; DMBT1, deleted in malignant brain tumors 1 protein; EEF1A1, elongation factor 1-alpha 1; GBP2, guanylate-binding protein 2; GBP6, guanylate-binding protein 6; GBP7, guanylate-binding protein 7; IFI6, interferon alpha-inducible protein 6; MBL1, mannose-binding protein A; SFTPD, pulmonary surfactant-associated protein D. **D.** Proteasome related to proteins with bat and human ranks. PSMA1, proteasome subunit alpha type-1; PSMA2, proteasome subunit alpha type-2; PSMA3, proteasome subunit alpha type-3; PSMA4, proteasome subunit alpha type-4; PSMA5, proteasome subunit alpha type-5; PSMA6, proteasome subunit alpha type-6; PSMA7, proteasome subunit alpha type-7; PSMB1, proteasome subunit beta type-1; PSMB2, proteasome subunit beta type-2; PSMB3, proteasome subunit beta type-3; PSMB4, proteasome subunit beta type-4; PSMB5, proteasome subunit beta type-5; PSMB6, proteasome subunit beta type-6; PSMD2, 26S proteasome non-ATPase regulatory subunit 2; RAD23A, UV excision repair protein RAD23 homolog A; RPS27A, ubiquitin-40S ribosomal protein S27a; USP14, ubiquitin carboxyl-terminal hydrolase 14. **E.** Redox related proteins with bat and human ranks. GPX1, glutathione peroxidase 1; GPX3, glutathione peroxidase 3; PRDX1, peroxiredoxin-1; PRDX2, peroxiredoxin-2; PRDX6, peroxiredoxin-6; RFESD, Rieske domain-containing protein; SOD1, superoxide dismutase [Cu-Zn]; TXN, thioredoxin; VNN2, vascular non-inflammatory molecule 2; VNN3, vascular noninflammatory molecule 3. **F.** Other uncategorized (“other”) proteins of interest with bat and human ranks. ACE, angiotensin-converting enzyme; ADIPOQ, adiponectin; AFP, alphafetoprotein; AHCY, adenosylhomocysteinase; CD177, CD177 antigen; CD97, CD97 antigen; CLCA1, calcium-activated chloride channel regulator 1; DPEP1, dipeptidase 1; EIF5A2, eukaryotic translation initiation factor 5A-2; FCGBP, IgGFc-binding protein; GCLM, glutamate--cysteine ligase regulatory subunit; HIST2H2BE, histone H2B type 2-E; HSPA2, heat shock-related 70 kDa protein 2; HSPA4, heat shock 70 kDa protein 4; HYAL1, hyaluronidase-1; IMPDH2, inosine-5’-monophosphate dehydrogenase 2; ITIH2, inter-alpha-trypsin inhibitor heavy chain H2; ITIH3, inter-alpha-trypsin inhibitor heavy chain H3; ITIH4, inter-alpha-trypsin inhibitor heavy chain H4; KLK1, kallikrein-1; LIPG, endothelial lipase; PAFAH1B2, platelet-activating factor acetylhydrolase IB subunit beta; PCBP1, poly(rC)-binding protein 1; PF4, platelet factor 4; SERPINB10, serpin B10; SMPD1, sphingomyelin phosphodiesterase; SRI, sorcin; TFRC, transferrin receptor protein 1; TMPRSS9, transmembrane protease serine 9. All protein ranks are given in Supplemental Table S5, while just select proteins here are described in Supplemental Table S6.

### Innate immunity

Investigation of apparent differences in proteins related to innate immunity is important because vampire bats harbor diverse pathogens ^22,51,54–56,74^ and because bats more generally appear to tolerate some viral infections without showing disease ^4,8,20,23^. In addition to the complement proteins described above, we identified notable proteins related to the innate immune system that are highly ranked in bat serum (Figure 2C). Given that some bats can constitutively express interferons (*e.g*., IFN-α in *Pteropus alecto*) ^18^, it was not surprising to detect interferon-inducible proteins (*e.g*., guanylate-binding proteins GBP2, GBP6, and GBP7). Of note is that IFN-α and IFN-γ induce many similar proteins in a highly dynamic manner ^75^. Specific to detected guanylate-binding proteins (GBPs), GBP2 was detected and shares antiviral properties with GBP5 (undetected), although circulating blood GBPs do not appear common in humans ^76–78^. It would be interesting to determine if certain proteins are stratified into exosomes, similar to the IFN-induced antiviral proteome contained in macrophage-derived exosomes ^79^. There were also proteins identified related to antiviral and antibacterial activity, notably relatively-high levels of CAMP (cathelicidin antimicrobial peptide). Finally, two scavenging proteins reported in human blood were ranked higher in bat serum: SFTPD (pulmonary surfactant-associated protein D) and DMBT1 (deleted in malignant brain tumors 1 protein). Though their functions are related, DMBT1 specifically binds viral and bacterial antigens ^80,81^ and can reduce viral infectivity ^82^. The question remains whether these proteins confer a unique ability to keep pathogen virulence at bay or if these elevated protein levels instead indicate an active infection. Ongoing studies will continue to define the host-pathogen relationship in species such as the vampire bat.

### Circulating proteasome

In addition to focusing on classically immunological proteins, we observed 14 proteasome subunits (Figure 2D). Specifically, 13 of the 14 known 20S subunits (though missing PSMB7) were observed along with one 26S component (PSMD2), indicating the presence of the circulating 20S proteasome ^83,84^. The lack of PSMB7 may be species-specific, similar to lower PSMB7 values reported across different mammal species ^85^. Although circulating 20S proteasome is lower ranked in the Human Blood Atlas, it has been detected in human plasma, possibly within exosomes ^86^. Given that our group has observed similarly abundant 20S proteasome subunits in other mammalian serum and plasma (*e.g*., California sea lion plasma proteome^87^), circulating 20S proteasome is likely present in vampire bats and not a technical artifact of hemolysis or serum preparation. The 20S proteasome mediates proteasomal cleavage, with numerous cellular roles including response to disease and oxidative stress ^84^. Interestingly, IFN-γ can induce formation of the immunoproteasome, which is the circulating 20S proteasome incorporated with β8, β9, and β10 subunits ^83^. The immunoproteasome has diverse functions, including cleaving peptides into MHC class I epitopes ^83^. The immunoproteasome is likely activated within hematopoietic cells, and may exist within extracellular vesicles though not necessarily circulating in the blood ^84^. Though these additional immunoproteasome subunits were not observed, it is interesting to hypothesize the role of such high levels of circulating 20S proteasome. These results demonstrate the potential of using mass spectrometry-based methods to study proteasomes, providing a basis for future immunological studies.

### Redox-related proteins

Beyond the possible uniqueness of bat immune systems is their ability to minimize or respond to oxidative stress. The evolution of flight in the chiropteran lineage was accompanied by mechanisms to minimize or repair the negative effects of oxidative stress generated by this metabolically costly activity, which may explain both the particular longevity of bats and their apparent viral tolerance ^8,88,89^. For reference in Figure 2C, we have included glutathione peroxidase levels, which do not appear very different from levels in humans, but we note that there are higher levels of other redox-related proteins in vampire bats. Increased levels of peroxiredoxins (PRDX1, PRDX2, and PRDX6), superoxide dismutase (SOD1), and thioredoxin (TXN) indicate an enhanced capacity to scavenge or oppose free-radicals. High-levels of Rieske domain-containing protein (RFESD), which facilitates binding excess iron, may be related to the iron-rich, blood diet of vampire bats, although ferritin light chain (FTL) levels were found to be similar between bats and humans, ranked 154 and 336, respectively. Finally, vascular noninflammatory molecule proteins (VNN2 and VNN3) have been described as being involved in redox reactions via the production of cysteamine ^90–92^. Overall, these results highlight that there may be increased oxidative resistance in vampire bats relative to humans; this pattern is likely to be conserved across bat species given the ubiquity of flight and long lifespans in bats, but this should be confirmed with broad comparative analyses.

### “Other” interesting proteins

In addition to proteins with obvious relation to known bat physiology, there are other interesting proteins that do not fit into a single category (Figure 2D). From this list, two sets of relationships are noteworthy: angiotensin-converting enzyme (ACE)/kallikrein 1 (KLK1) and hyaluronan-related proteins. Both ACE and KLK1 are higher in bats than humans, though their functions are antagonistic: kallikrein increases bradykinin, which vasodilates, while ACE cleaves bradykinin and thus may reduce vasodilation. In comparison, angiotensinogen (AGT), a source of the vasoconstrictor peptide angiotensin 2, is almost the same rank in bats as humans (ranked 29 and 25, respectively; Supplemental Table S5). Of note, given the entry of SARS-CoV-2 in humans, is the detection of TMPRSS9 as a mid-abundance protein in bat serum, though ACE2, which is detected in humans (rank 2517; Supplemental Table S5), was not present in bat serum. There are also increased hyaluronan-related proteins in vampire bat serum. The hyaluronan receptor CD44 is the same rank in bats and humans (Supplemental Table S5), though two hyaluronan carrier proteins (ITIH2 and ITIH3) and hyaluronidase 1 (HYAL1), which hydrolyzes hyaluronan, are higher in bats (Figure 2F). This last protein, HYAL1, is ranked 3024 in human plasma, versus 173 in vampire bat serum. It is unclear whether this indicates high levels of hyaluronan in bat serum, which in the naked-mole rat may confer anti-cancer properties ^93^, or that these proteins are involved with keeping serum hyaluronan levels low in response to infection ^94^. There are additional undiscussed proteins listed in this “other” category, as well as a full list of protein ranks compared to the Human Blood Atlas (Supplemental Table S5-S6), that researchers are encouraged to browse for insight beyond the survey nature of this manuscript. It is interesting to postulate that vampire bat blood may have an anti-pathogenic phenotype, but it is unclear if this is in response to infection or constitutive. Similarly, given that many antiviral proteins are often involved in anti-cancer roles ^95^, these two roles could plausibly be connected.

### Pathogen detection

Data-independent acquisition (DIA) is an emerging technique in mass spectrometry-based proteomics, which in theory should allow deeper coverage than typical data-dependent acquisition (DDA)-based analysis by overcoming the stochastic nature of DDA data. We did not set out to compare the two approaches, but found this to be an interesting use-case of DIA to analyze undepleted serum. Detecting 361 proteins in undepleted serum using a relatively short 60 min gradient demonstrated that DIA-based proteomics of undepleted serum can scale reasonably well. Moreover, we took advantage of this technique to evaluate the presence of viral proteins. *Desmodus rotundus* has tested positive elsewhere in its geographic range for adenoviruses, coronaviruses, flaviviruses, hantaviruses, herpesviruses, and paramyxoviruses, as well as for antibodies against henipavirus-like viruses ^51,54–62^.Searches were performed with *Rabies lyssavirus*, Adenoviridae, *Betacoronavirus*, *Henipavirus*, Herpesvirales, and *Morbillivirus* databases. Using this approach, we detected no viral proteins in the human pooled sample but identified two viral peptides in the bat sera (Supplemental Figures 2–3): PRSGIPDR from the Rh186 protein in *Macacine herpesvirus 3* (i.e., Rhesus cytomegalovirus) and LVTTEVK from the ORF1a protein in Middle East respiratory syndrome-related coronavirus (MERS-CoV). Detection of the MERS-CoV protein is of particular interest, as other betacoronaviruses have been detected in vampire bats and cell line studies suggest their receptors (i.e., DPP4) can support MERS-CoV replication ^59,96^. Although we are unaware of specific MERS-CoV detection in vampire bats, a betacoronavirus with high amino acid similarity to MERS-CoV has been identified in an unrelated but sympatric bat species in Mexico (*Nyctinomops laticaudatus*) ^97^. More generally, these peptides were detected using a relatively short gradient separation of undepleted serum, suggesting that a method enriching for viral particles prior to analysis could easily detect more viral proteins if present. Importantly, although PCR-based techniques are frequently used for viral surveillance in bats, a mass spectrometry-based approach could query a larger search space while maintaining acceptable detection limits. In cases where a specific viral taxon was targeted, mass spectrometry could provide a more accurate method of virus detection (e.g., as evidenced by mass spectrometry-based SARS-CoV-2 detection in humans ^98–101^).

Targeted mass spectrometry methods using stable isotope-labeled peptides allow for creation of sensitive, precise, and accurate assays that avoid interferences that plague antibody-based methods. The first step of developing these targeted assays is empirically confirming the presence of predicted peptides, which is possible even in non-model species. For example, shotgun proteomics were first used to identify proteotypic adiponectin peptides in dolphin plasma, followed by validation of a parallel-reaction monitoring method to measure adiponectin at nmol/L levels using heavy isotope labeled peptides ^102^. Any of the proteins identified in the current study can be measured using this technique. For example, PRM assays could be constructed for each of the 31 complement activation proteins listed, and this could be performed in high-throughput manner. These assays would be free from measurement interference (*i.e*., cross-reactivity seen in antibody assays) with all the benefits of accuracy and precision found in PRM-based mass spectrometry assays ^103,104^.

## Conclusion

Our results highlight the power of modern proteomic techniques to provide new insights where other techniques might struggle. There is tremendous value in genomic and transcriptomic analyses; however, to understand the molecular landscape of blood, proteomic analysis is unique. This survey of the serum proteome of a small group of vampire bats provides a first look at the abundance of immunological proteins as well as proteins with unclear roles in the vampire bat phenotype. Moreover, we show that mass spectrometry-based viral detection in serum is possible and may be a viable tool for pathogen surveillance. Overall, these data can be used to develop targeted assays for future vampire bat research while also serving as a demonstration of the potential of proteomic studies for research on various topics in other bat species. For example, the physiology of white-nose syndrome and longevity are important topics that could benefit from investigations using modern biomolecular analytical techniques like proteomics. Future work will continue to investigate if proteins elevated in vampire bats compared to humans are similarly elevated in other mammals, or whether they are more common in wild animals than expected and not specific to a vampire bat phenotype. There is an ongoing need for comparative physiological studies to move beyond two-species comparisons and broaden our horizons across multiple clades of the tree of life.

## Supplemental Tables

Supplemental Table S1 Experimental key with sample metadata

Supplemental Table S2 Intermediate data table of vampire bat serum protein ortholog mapping and quantities

Supplemental Table S3 Vampire bat serum protein orthologs and quantities

Supplemental Table S4 NIST 909c human serum protein relative quantities

Supplemental Table S5 Rank comparisons of the vampire bat proteome to the Human Blood Atlas

Supplemental Table S6 Proteins of interest with rank comparisons between vampire bat proteome and the Human Blood Atlas

## Supplemental Figures

Supplemental Figure 1 Relative standard deviation distribution of protein abundance in technical replicate

Supplemental Figure 2 Rh186 viral peptide identification

Supplemental Figure 3 ORF1a viral peptide identification

## Acknowledgements

For assistance with bat sampling logistics and research permits, we thank Brock Fenton, Mark Howells, Neil Duncan, and staff of the Lamanai Field Research Center. Identification of certain commercial equipment, instruments, software or materials does not imply recommendation or endorsement by the National Institute of Standards and Technology, nor does it imply that the products identified are necessarily the best available for the purpose.

---

**Supplemental Figure S1.**
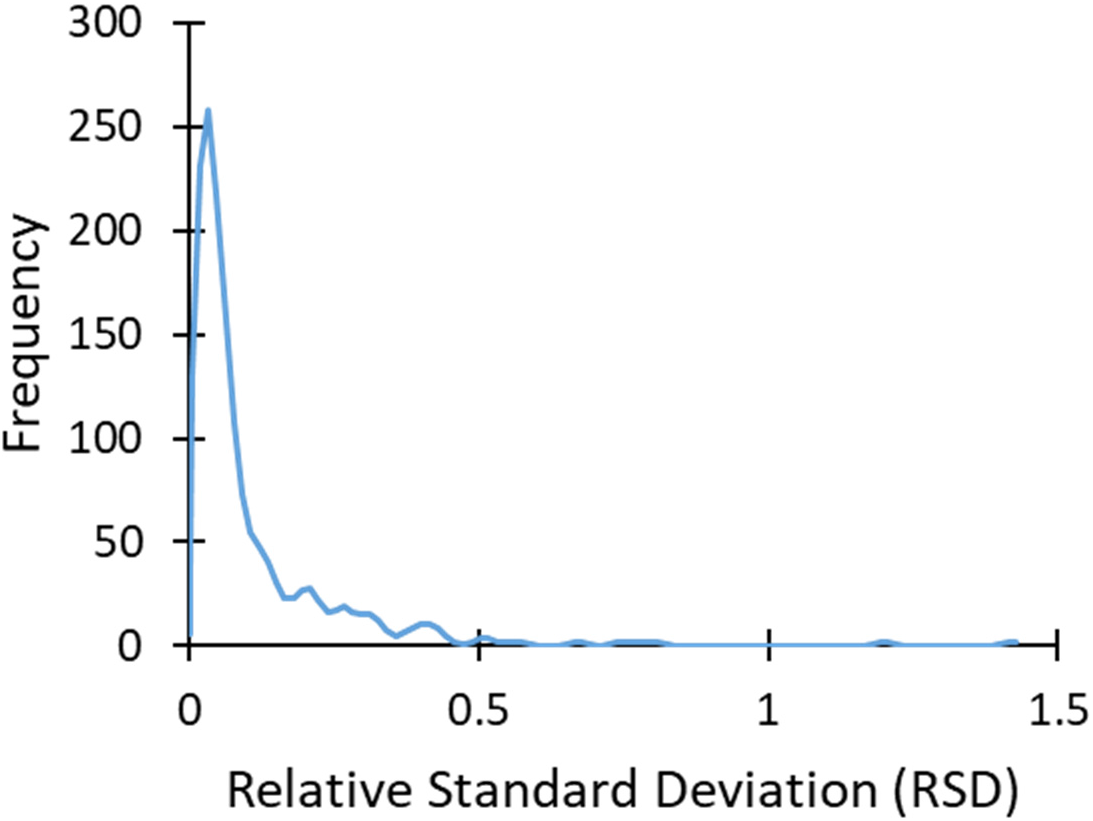
Distribution of relative standard deviations in technical replicate. One sample was injected in triplicate to evaluate technical performance of the method. Of the 361 proteins detected across the experiment, between 25 and 30 were filtered out, in total only 322 proteins were identified in all three injections. Using relative abundance from these 322 proteins, relative standard deviation (RSD) was calculated. The average RSD was 10.6 %, median RSD was 5.1 %, and 270 (83.9 % of 322) had an RSD < 20 %.

**Supplemental Figure S2.**
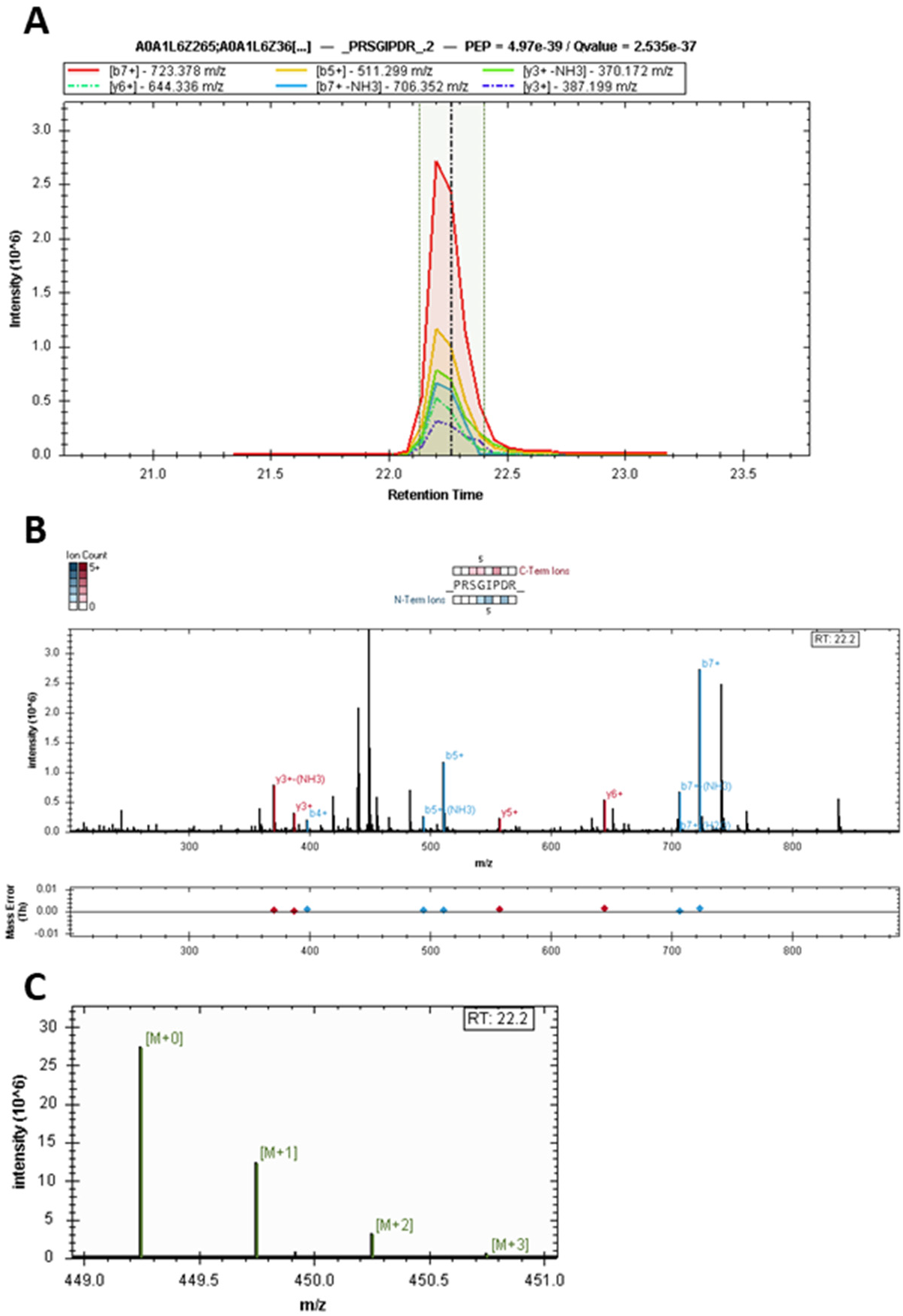
Rh186 viral peptide identification. **A.** Fragmentation ions matching to PRSGIPDR from Rh186 protein in Macacine herpesvirus 3 (i.e., Rhesus cytomegalovirus); UniProt IDs A0A1L6Z265; A0A1L6Z360; I3WF26; Q2FAB1; Q7TFF6. **B.** Matched b- and y-ions are shown along with mass errors. **C.** Precursor isotope pattern. Spectra are from Bat_14.

**Supplemental Figure S3.**
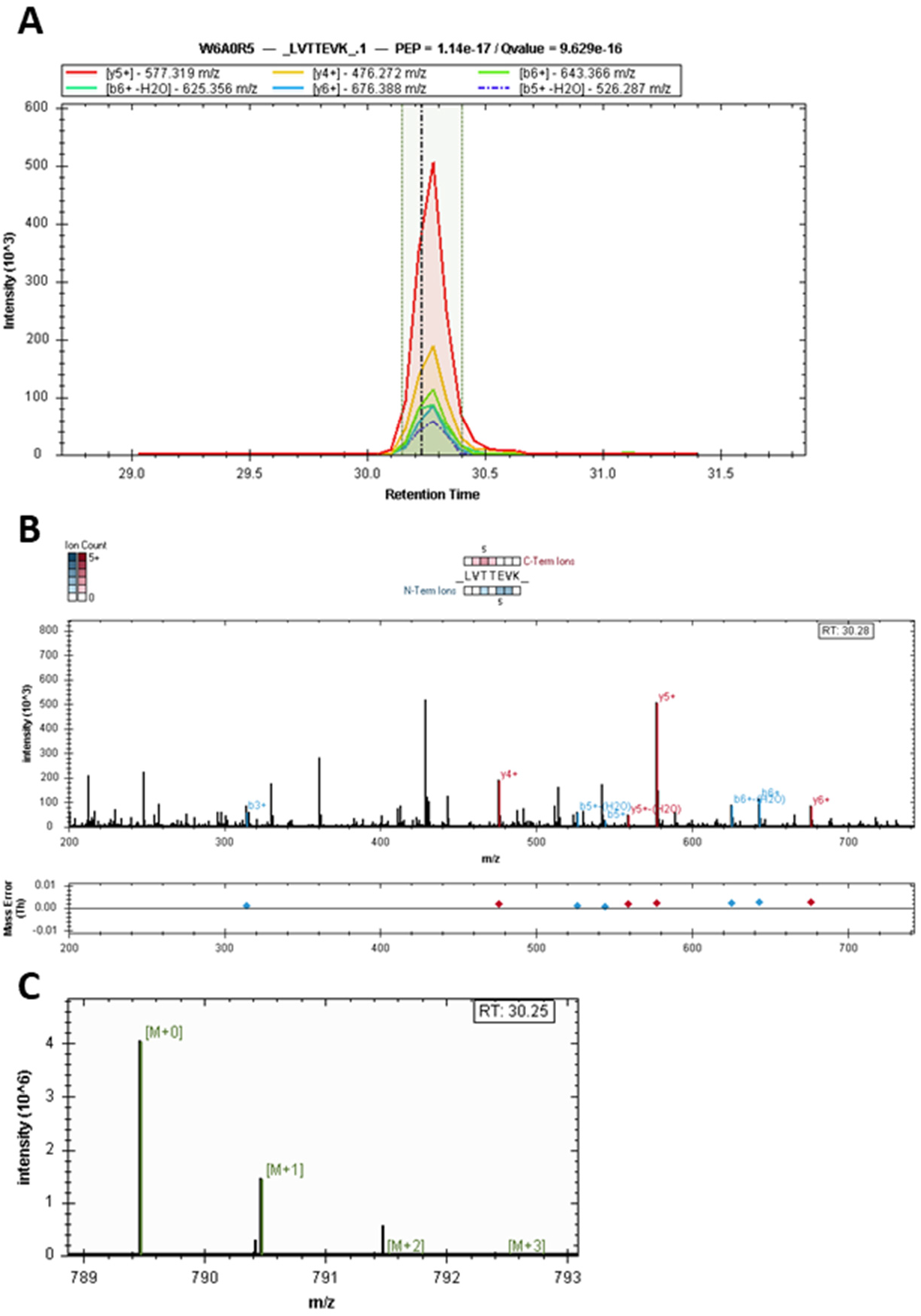
ORF1a viral peptide identification. **A.** Fragmentation ions matching to LVTTEVK from Middle East respiratory syndrome-related coronavirus (MERS-CoV), UniProt ID W6A0R5. **B.** Matched b- and y-ions are shown along with mass errors. **C.** Precursor isotope pattern. Spectra are from Bat_14.

## References

(1) Wilson, D.E.; Reeder, D. M. Mammal Species of the World: A Taxonomic and Geographic Reference; JHU Press, 2005.

(2) Kunz, T. H.; Braun de Torrez, E.; Bauer, D.; Lobova, T.; Fleming, T. H. Ecosystem Services Provided by Bats. Ann. N. Y. Acad. Sci. 2011, 1223, 1–38.

(3) Wilkinson, G. S.; South, J. M. Life History, Ecology and Longevity in Bats. Aging Cell 2002, 1 (2), 124–131.

(4) Jebb, D.; Huang, Z.; Pippel, M.; Hughes, G. M.; Lavrichenko, K.; Devanna, P.; Winkler, S.; Jermiin, L. S.; Skirmuntt, E. C.; Katzourakis, A.; Burkitt-Gray, L.; Ray, D. A.; Sullivan, K. A. M.; Roscito, J. G.; Kirilenko, B. M.; Dávalos, L. M.; Corthals, A. P.; Power, M. L.; Jones, G.; Ransome, R. D.; Dechmann, D. K. N.; Locatelli, A. G.; Puechmaille, S. J.; Fedrigo, O.; Jarvis, E. D.; Hiller, M.; Vernes, S. C.; Myers, E. W.; Teeling, E. C. Six Reference-Quality Genomes Reveal Evolution of Bat Adaptations. Nature 2020, 583 (7817), 578–584.

(5) Ingala, M. R.; Simmons, N. B.; Perkins, S. L. Bats Are an Untapped System for Understanding Microbiome Evolution in Mammals. mSphere 2018, 3 (5). https://doi.org/10.1128/mSphere.00397-18.

(6) Ripperger, S. P.; Carter, G. G.; Duda, N.; Koelpin, A.; Cassens, B.; Kapitza, R.; Josic, D.; Berrío-Martínez, J.; Page, R. A.; Mayer, F. Vampire Bats That Cooperate in the Lab Maintain Their Social Networks in the Wild. Curr. Biol. 2019, 29 (23), 4139–4144.e4.

(7) Guth, S.; Visher, E.; Boots, M.; Brook, C.E. Host Phylogenetic Distance Drives Trends in Virus Virulence and Transmissibility across the Animal–human Interface. Philos. Trans. R. Soc. Lond. B Biol. Sci. 2019, 374 (1782), 20190296.

(8) Brook, C. E.; Dobson, A. P. Bats as “special”reservoirs for Emerging Zoonotic Pathogens. Trends Microbiol. 2015, 23 (3), 172–180.

(9) Olival, K. J.; Hosseini, P. R.; Zambrana-Torrelio, C.; Ross, N.; Bogich, T. L.; Daszak, P. Host and Viral Traits Predict Zoonotic Spillover from Mammals. Nature 2017, 546 (7660), 646–650.

(10) Halpin, K.; Hyatt, A. D.; Fogarty, R.; Middleton, D.; Bingham, J.; Epstein, J. H.; Rahman, S. A.; Hughes, T.; Smith, C.; Field, H. E.; Daszak, P.; Henipavirus Ecology Research Group. Pteropid Bats Are Confirmed as the Reservoir Hosts of Henipaviruses: A Comprehensive Experimental Study of Virus Transmission. Am. J. Trop. Med. Hyg. 2011, 85 (5), 946–951.

(11) Amman, B. R.; Jones, M. E. B.; Sealy, T. K.; Uebelhoer, L. S.; Schuh, A. J.; Bird, B. H.; Coleman-McCray, J. D.; Martin, B. E.; Nichol, S. T.; Towner, J. S. Oral Shedding of Marburg Virus in Experimentally Infected Egyptian Fruit Bats (Rousettus Aegyptiacus). J. Wildl. Dis. 2015, 51 (1), 113–124.

(12) Banyard, A. C.; Hayman, D.; Johnson, N.; McElhinney, L.; Fooks, A. R. Bats and Lyssaviruses. Adv. Virus Res. 2011, 79, 239–289.

(13) Li, W.; Shi, Z.; Yu, M.; Ren, W.; Smith, C.; Epstein, J. H.; Wang, H.; Crameri, G.; Hu, Z.; Zhang, H.; Zhang, J.; McEachern, J.; Field, H.; Daszak, P.; Eaton, B. T.; Zhang, S.; Wang, L.-F. Bats Are Natural Reservoirs of SARS-like Coronaviruses. Science 2005, 310 (5748), 676–679.

(14) Plowright, R. K.; Eby, P.; Hudson, P. J.; Smith, I. L.; Westcott, D.; Bryden, W. L.; Middleton, D.; Reid, P. A.; McFarlane, R. A.; Martin, G.; Tabor, G. M.; Skerratt, L. F.; Anderson, D. L.; Crameri, G.; Quammen, D.; Jordan, D.; Freeman, P.; Wang, L.-F.; Epstein, J. H.; Marsh, G. A.; Kung, N. Y.; McCallum, H. Ecological Dynamics of Emerging Bat Virus Spillover. Proc. Biol. Sci. 2015, 282 (1798), 20142124.

(15) Kessler, M. K.; Becker, D. J.; Peel, A. J. Changing Resource Landscapes and Spillover of Henipaviruses. Ann. N. Y. Acad. Sci. 2018.

(16) Williamson, M. M.; Hooper, P. T.; Selleck, P. W.; Westbury, H. A.; Slocombe, R. F. Experimental Hendra Virus Infectionin Pregnant Guinea-Pigs and Fruit Bats (Pteropus Poliocephalus). J. Comp. Pathol. 2000, 122 (2-3), 201–207.

(17) Mollentze, N.; Streicker, D. G. Viral Zoonotic Risk Is Homogenous among Taxonomic Orders of Mammalian and Avian Reservoir Hosts. Proceedings of the National Academy of Sciences. 2020, pp 9423–9430. https://doi.org/10.1073/pnas.1919176117.

(18) Zhou, P.; Tachedjian, M.; Wynne, J. W.; Boyd, V.; Cui, J.; Smith, I.; Cowled, C.; Ng, J. H. J.; Mok, L.; Michalski, W. P.; Mendenhall, I. H.; Tachedjian, G.; Wang, L.-F.; Baker, M. L. Contraction of the Type I IFN Locus and Unusual Constitutive Expression of IFN-α in Bats. Proc. Natl. Acad. Sci. U. S. A. 2016, 113 (10), 2696–2701.

(19) Becker, D. J.; Czirják, G. Á. ; Rynda-Apple, A.; Plowright, R. K. Handling Stress and Sample Storage Are Associated with Weaker Complement-Mediated Bactericidal Ability in Birds but Not Bats. Physiological and Biochemical Zoology. 2019, pp 37–48. https://doi.org/10.1086/701069.

(20) Schountz, T.; Baker, M. L.; Butler, J.; Munster, V. Immunological Control of Viral Infections in Bats and the Emergence of Viruses Highly Pathogenic to Humans. Front. Immunol. 2017, 8, 1098.

(21) Becker, D. J.; Crowley, D. E.; Washburne, A. D.; Plowright, R. K. Temporal and Spatial Limitations in Global Surveillance for Bat Filoviruses and Henipaviruses. Biol. Lett. 2019, 15 (12), 20190423.

(22) Bergner, L. M.; Orton, R. J.; Benavides, J. A.; Becker, D. J.; Tello, C.; Biek, R.; Streicker, D. G. Demographic and Environmental Drivers of Metagenomic Viral Diversity in Vampire Bats. Mol. Ecol. 2020, 29 (1), 26–39.

(23) Brook, C. E.; Boots, M.; Chandran, K.; Dobson, A. P.; Drosten, C.; Graham, A. L.; Grenfell, B. T.; Müller, M. A.; Ng, M.; Wang, L.-F.; van Leeuwen, A. Accelerated Viral Dynamics in Bat Cell Lines, with Implications for Zoonotic Emergence. Elife 2020, 9. https://doi.org/10.7554/eLife.48401.

(24) Sikes, R. S.; Gannon, W. L. Guidelines of the American Society of Mammalogists for the Use of Wild Mammals in Research. J. Mammal. 2011, 92 (1), 235–253.

(25) Jones, K. E.; Bielby, J.; Cardillo, M.; Fritz, S. A.; O’Dell, J.; Orme, C. D. L.; Safi, K.; Sechrest, W.; Boakes, E. H.; Carbone, C.; Connolly, C.; Cutts, M. J.; Foster, J. K.; Grenyer, R.; Habib, M.; Plaster, C. A.; Price, S. A.; Rigby, E. A.; Rist, J.; Teacher, A.; Bininda-Emonds, O. R. P.; Gittleman, J. L.; Mace, G. M.; Purvis, A. PanTHERIA: A Species-Level Database of Life History, Ecology, and Geography of Extant and Recently Extinct Mammals. Ecology. 2009, pp 2648–2648. https://doi.org/10.1890/08-1494.1.

(26) Baker, M. L.; Schountz, T.; Wang, L.-F. Antiviral Immune Responses of Bats: A Review. Zoonoses Public Health 2013, 60 (1), 104–116.

(27) Schountz, T. Immunology of Bats and Their Viruses: Challenges and Opportunities. Viruses 2014, 6 (12), 4880–4901.

(28) Allen, L. C.; Turmelle, A. S.; Mendonça, M. T.; Navara, K. J.; Kunz, T. H.; McCracken, G. F. Roosting Ecology and Variation in Adaptive and Innate Immune System Function in the Brazilian Free-Tailed Bat (Tadarida Brasiliensis). J. Comp. Physiol. B 2009, 179 (3), 315–323.

(29) Becker, D. J.; Czirják, G. Á.; Volokhov, D. V. Livestock Abundance Predicts Vampire Bat Demography, Immune Profiles and Bacterial Infection Risk. of the Royal… 2018.

(30) Ruoss, S.; Becker, N. I.; Otto, M. S.; Czirják, G. Á.; Encarnação, J. A. Effect of Sex and Reproductive Status on the Immunity of the Temperate Bat Myotis Daubentonii. Mamm. Biol. 2019, 94 (1), 120–126.

(31) Young, C. C. W.; Olival, K. J. Optimizing Viral Discovery in Bats. PLoS One 2016, 11 (2), e0149237.

(32) Stepanian, P. M.; Wainwright, C. E. Ongoing Changes in Migration Phenology and Winter Residency at Bracken Bat Cave. Glob. Chang. Biol. 2018, 24 (7), 3266–3275.

(33) Plowright, R. K.; Peel, A. J.; Streicker, D. G.; Gilbert, A. T.; McCallum, H.; Wood, J.; Baker, M. L.; Restif, O. Transmission or Within-Host Dynamics Driving Pulses of Zoonotic Viruses in Reservoir-Host Populations. PLoS Negl. Trop. Dis. 2016, 10 (8), e0004796.

(34) Gilbert, A. T.; Fooks, A. R.; Hayman, D. T. S.; Horton, D. L.; Müller, T.; Plowright, R.; Peel, A. J.; Bowen, R.; Wood, J. L. N.; Mills, J.; Cunningham, A. A.; Rupprecht, C. E. Deciphering Serology to Understand the Ecology of Infectious Diseases in Wildlife. Ecohealth 2013, 10 (3), 298–313.

(35) Gerrard, D. L.; Hawkinson, A.; Sherman, T.; Modahl, C. M.; Hume, G.; Campbell, C. L.; Schountz, T.; Frietze, S. Transcriptomic Signatures of Tacaribe Virus-Infected Jamaican Fruit Bats. mSphere 2017, 2 (5). https://doi.org/10.1128/mSphere.00245-17.

(36) Lee, A. K.; Kulcsar, K. A.; Elliott, O.; Khiabanian, H.; Nagle, E. R.; Jones, M. E. B.; Amman, B. R.; Sanchez-Lockhart, M.; Towner, J. S.; Palacios, G.; Rabadan, R. De Novo Transcriptome Reconstruction and Annotation of the Egyptian Rousette Bat. BMC Genomics 2015, 16, 1033.

(37) Bergner, L. M.; Orton, R. J.; da Silva Filipe, A.; Shaw, A. E.; Becker, D. J.; Tello, C.; Biek, R.; Streicker, D. G. Using Noninvasive Metagenomics to Characterize Viral Communities from Wildlife. Mol. Ecol. Resour. 2019, 19 (1), 128–143.

(38) Donaldson, E. F.; Haskew, A. N.; Gates, J. E.; Huynh, J.; Moore, C. J.; Frieman, M. B. Metagenomic Analysis of the Viromes of Three North American Bat Species: Viral Diversity among Different Bat Species That Share a Common Habitat. J. Virol. 2010, 84 (24), 13004–13018.

(39) Uhlén, M.; Karlsson, M. J.; Hober, A.; Svensson, A.-S.; Scheffel, J.; Kotol, D.; Zhong, W.; Tebani, A.; Strandberg, L.; Edfors, F.; Sjöstedt, E.; Mulder, J.; Mardinoglu, A.; Berling, A.; Ekblad, S.; Dannemeyer, M.; Kanje, S.; Rockberg, J.; Lundqvist, M.; Malm, M.; Volk, A.-L.; Nilsson, P.; Månberg, A.; Dodig-Crnkovic, T.; Pin, E.; Zwahlen, M.; Oksvold, P.; von Feilitzen, K.; Häussler, R. S.; Hong, M.-G.; Lindskog, C.; Ponten, F.; Katona, B.; Vuu, J.; Lindström, E.; Nielsen, J.; Robinson, J.; Ayoglu, B.; Mahdessian, D.; Sullivan, D.; Thul, P.; Danielsson, F.; Stadler, C.; Lundberg, E.; Bergström, G.; Gummesson, A.; Voldborg, B. G.; Tegel, H.; Hober, S.; Forsström, B.; Schwenk, J. M.; Fagerberg, L.; Sivertsson, Å. The Human Secretome. Sci. Signal. 2019, 12 (609). https://doi.org/10.1126/scisignal.aaz0274.

(40) Geyer, P. E.; Holdt, L. M.; Teupser, D.; Mann, M. Revisiting Biomarker Discovery by Plasma Proteomics. Mol. Syst. Biol. 2017, 13 (9), 942.

(41) Hecht, A. M.; Braun, B. C.; Krause, E.; Voigt, C. C.; Greenwood, A. D.; Czirják, G. Á. Plasma Proteomic Analysis of Active and Torpid Greater Mouse-Eared Bats (Myotis Myotis). Sci. Rep. 2015, 5, 16604.

(42) Hecht-Höger, A. M.; Braun, B. C.; Krause, E.; Meschede, A.; Krahe, R.; Voigt, C. C.; Greenwood, A. D.; Czirják, G. Á. Plasma Proteomic Profiles Differ between European and North American Myotid Bats Colonized by Pseudogymnoascus Destructans. Mol. Ecol. 2020, 29 (9), 1745–1755.

(43) Francischetti, I. M. B.; Assumpção, T. C. F.; Ma, D.; Li, Y.; Vicente, E. C.; Uieda, W.; Ribeiro, J. M. C. The “Vampirome”: Transcriptome and Proteome Analysis of the Principal and Accessory Submaxillary Glands of the Vampire Bat Desmodus Rotundus, a Vector of Human Rabies. J. Proteomics 2013, 82, 288–319.

(44) Yin, Q.; Zhang, Y.; Dong, D.; Lei, M.; Zhang, S.; Liao, C.-C.; Pan, Y.-H. Maintenance of Neural Activities in Torpid Rhinolophus Ferrumequinum Bats Revealed by 2D Gel-Based Proteome Analysis. Biochim. Biophys. Acta: Proteins Proteomics 2017, 1865 (8), 1004–1019.

(45) Woon, A. P.; Boyd, V.; Todd, S.; Smith, I.; Klein, R.; Woodhouse, I. B.; Riddell, S.; Crameri, G.; Bingham, J.; Wang, L.-F.; Purcell, A. W.; Middleton, D.; Baker, M. L. Acute Experimental Infection of Bats and Ferrets with Hendra Virus: Insights into the Early Host Response of the Reservoir Host and Susceptible Model Species. PLoS Pathog. 2020, 16 (3), e1008412.

(46) Greenhall, A. M. Natural History of Vampire Bats; CRC Press, 2018.

(47) Bobrowiec, P. E. D.; Lemes, M. R.; Gribel, R. Prey Preference of the Common Vampire Bat (Desmodus Rotundus, Chiroptera) Using Molecular Analysis. J. Mammal. 2015, 96 (1), 54–63.

(48) Streicker, D. G.; Allgeier, J. E. Foraging Choices of Vampire Bats in Diverse Landscapes: Potential Implications for Land-Use Change and Disease Transmission. J. Appl. Ecol. 2016, 53 (4), 1280–1288.

(49) Schneider, M. C.; Romijn, P. C.; Uieda, W.; Tamayo, H.; da Silva, D. F.; Belotto, A.; da Silva, J. B.; Leanes, L. F. Rabies Transmitted by Vampire Bats to Humans: An Emerging Zoonotic Disease in Latin America? Rev. Panam. Salud Publica 2009, 25 (3), 260–269.

(50) Condori-Condori, R. E.; Streicker, D. G.; Cabezas-Sanchez, C.; Velasco-Villa, A. Enzootic and Epizootic Rabies Associated with Vampire Bats, Peru. Emerg. Infect. Dis. 2013, 19 (9). https://doi.org/10.3201/eid1809.130083.

(51) Wray, A. K.; Olival, K. J.; Morán, D.; Lopez, M. R.; Alvarez, D.; Navarrete-Macias, I.; Liang, E.; Simmons, N. B.; Lipkin, W. I.; Daszak, P.; Anthony, S. J. Viral Diversity, Prey Preference, and Bartonella Prevalence in Desmodus Rotundus in Guatemala. Ecohealth 2016, 13 (4), 761–774.

(52) Patterson, C. Deforestation, Agricultural Intensification, and Farm Resilience in Northern Belize: 1980–2010, University of Otago, 2016.

(53) Becker, D. J.; Broos, A.; Bergner, L. M.; Meza, D. K.; Simmons, N. B.; Brock Fenton, M.; Altizer, S.; Streicker, D. G. Temporal Patterns of Vampire Bat Rabies and Host Connectivity in Belize. bioRxiv, 2020, 2020.07.16.204446. https://doi.org/10.1101/2020.07.16.204446.

(54) Drexler, J. F.; Corman, V. M.; Müller, M. A.; Maganga, G. D.; Vallo, P.; Binger, T.; Gloza-Rausch, F.; Cottontail, V. M.; Rasche, A.; Yordanov, S.; Seebens, A.; Knörnschild, M.; Oppong, S.; Adu Sarkodie, Y.; Pongombo, C.; Lukashev, A. N.; Schmidt-Chanasit, J.; Stöcker, A.; Carneiro, A. J. B.; Erbar, S.; Maisner, A.; Fronhoffs, F.; Buettner, R.; Kalko, E. K. V.; Kruppa, T.; Franke, C. R.; Kallies, R.; Yandoko, E. R. N.; Herrler, G.; Reusken, C.; Hassanin, A.; Krüger, D. H.; Matthee, S.; Ulrich, R. G.; Leroy, E. M.; Drosten, C. Bats Host Major Mammalian Paramyxoviruses. Nat. Commun. 2012, 3, 796.

(55) Escalera-Zamudio, M.; Taboada, B.; Rojas-Anaya, E.; Löber, U.; Loza-Rubio, E.; Arias, C. F.; Greenwood, A. D. Viral Communities Among Sympatric Vampire Bats and Cattle. Ecohealth 2018, 15 (1), 132–142.

(56) de Araujo, J.; Lo, M. K.; Tamin, A.; Ometto, T. L.; Thomazelli, L. M.; Nardi, M. S.; Hurtado, R. F.; Nava, A.; Spiropoulou, C. F.; Rota, P. A.; Durigon, E. L. Antibodies Against Henipa-Like Viruses in Brazilian Bats. Vector Borne Zoonotic Dis. 2017, 17 (4), 271–274.

(57) Bergner, L. M.; Orton, R. J.; Broos, A.; Tello, C.; Becker, D. J.; Carrera, J. E.; Patel, A. H.; Biek, R.; Streicker, D. G. Diversification of Mammalian Deltaviruses by Host Shifting, 2020, 2020.06.17.156745. https://doi.org/10.1101/2020.06.17.156745.

(58) Asano, K. M.; Hora, A. S.; Scheffer, K. C.; Fahl, W. O.; Iamamoto, K.; Mori, E.; Brandão, P. E. Erratum to: Alphacoronavirus in Urban Molossidae and Phyllostomidae Bats, Brazil. Virol. J. 2016, 13 (1), 124.

(59) Brandão, P. E.; Scheffer, K.; Villarreal, L. Y.; Achkar, S.; Oliveira, R. de N.; Fahl, W. de O.; Castilho, J. G.; Kotait, I.; Richtzenhain, L. J. A Coronavirus Detected in the Vampire Bat Desmodus Rotundus. Braz. J. Infect. Dis. 2008, 12 (6), 466–468.

(60) Quan, P.-L.; Firth, C.; Conte, J. M.; Williams, S. H.; Zambrana-Torrelio, C. M.; Anthony, S. J.; Ellison, J. A.; Gilbert, A. T.; Kuzmin, I. V.; Niezgoda, M.; Osinubi, M. O. V.; Recuenco, S.; Markotter, W.; Breiman, R. F.; Kalemba, L.; Malekani, J.; Lindblade, K. A.; Rostal, M. K.; Ojeda-Flores, R.; Suzan, G.; Davis, L. B.; Blau, D. M.; Ogunkoya, A. B.; Alvarez Castillo, D. A.; Moran, D.; Ngam, S.; Akaibe, D.; Agwanda, B.; Briese, T.; Epstein, J. H.; Daszak, P.; Rupprecht, C. E.; Holmes, E. C.; Lipkin, W. I. Bats Are a Major Natural Reservoir for Hepaciviruses and Pegiviruses. Proc. Natl. Acad. Sci. U. S. A. 2013, 110 (20), 8194–8199.

(61) James, S.; Donato, D.; de Thoisy, B.; Lavergne, A.; Lacoste, V. Novel Herpesviruses in Neotropical Bats and Their Relationship with Other Members of the Herpesviridae Family. Infect. Genet. Evol. 2020, 84, 104367.

(62) Chen, L.; Liu, B.; Yang, J.; Jin, Q. DBatVir: The Database of Bat-Associated Viruses. Database 2014, 2014, bau021.

(63) Delpietro, H. A.; Russo, R. G. Observations of the Common Vampire Bat (Desmodus Rotundus) and the Hairy-Legged Vampire Bat (Diphylla Ecaudata) in Captivity. Mamm. Biol. 2002, 67 (2), 65–78.

(64) Perez-Riverol, Y.; Csordas, A.; Bai, J.; Bernal-Llinares, M.; Hewapathirana, S.; Kundu, D. J.; Inuganti, A.; Griss, J.; Mayer, G.; Eisenacher, M.; Pérez, E.; Uszkoreit, J.; Pfeuffer, J.; Sachsenberg, T.; Yilmaz, S.; Tiwary, S.; Cox, J.; Audain, E.; Walzer, M.; Jarnuczak, A. F.; Ternent, T.; Brazma, A.; Vizcaíno, J. A. The PRIDE Database and Related Tools and Resources in 2019: Improving Support for Quantification Data. Nucleic Acids Res. 2019, 47 (D1), D442–D450.

(65) Camacho, C.; Coulouris, G.; Avagyan, V.; Ma, N.; Papadopoulos, J.; Bealer, K.; Madden, T. L. BLAST+: Architecture and Applications. BMC Bioinformatics 2009, 10, 421.

(66) Dunkelberger, J. R.; Song, W.-C. Complement and Its Role in Innate and Adaptive Immune Responses. Cell Res. 2010, 20 (1), 34–50.

(67) Nordahl, E. A.; Rydengård, V.; Nyberg, P.; Nitsche, D. P.; Mörgelin, M.; Malmsten, M.; Björck, L.; Schmidtchen, A. Activation of the Complement System Generates Antibacterial Peptides. Proc. Natl. Acad. Sci. U. S. A. 2004, 101 (48), 16879–16884.

(68) Ricklin, D.; Hajishengallis, G.; Yang, K.; Lambris, J. D. Complement: A Key System for Immune Surveillance and Homeostasis. Nat. Immunol. 2010, 11 (9), 785–797.

(69) Moore, M. S.; Reichard, J. D.; Murtha, T. D.; Zahedi, B.; Fallier, R. M.; Kunz, T. H. Specific Alterations in Complement Protein Activity of Little Brown Myotis (Myotis Lucifugus) Hibernating in White-Nose Syndrome Affected Sites. PLoS One 2011, 6 (11), e27430.

(70) Pilosof, S.; Korine, C.; Moore, M. S.; Krasnov, B. R. Effects of Sewage-Water Contamination on the Immune Response of a Desert Bat. Mamm. Biol. 2014, 79 (3), 183–188.

(71) Anderson, N. L.; Anderson, N. G. The Human Plasma Proteome: History, Character, and Diagnostic Prospects. Mol. Cell. Proteomics 2002, 1 (11), 845–867.

(72) The human blood in proteins detected in ms - The Human Protein Atlas https://www.proteinatlas.org/humanproteome/blood/proteins+detected+in+ms (accessed Jul 19, 2020).

(73) Geyer, P. E.; Voytik, E.; Treit, P. V.; Doll, S.; Kleinhempel, A.; Niu, L.; Müller, J. B.; Buchholtz, M.-L.; Bader, J. M.; Teupser, D.; Holdt, L. M.; Mann, M. Plasma Proteome Profiling to Detect and Avoid Sample-Related Biases in Biomarker Studies. EMBO Mol. Med. 2019, 11 (11), e10427.

(74) Correa-Giron, P.; Calisher, C. H.; Baer, G. M. Epidemic Strain of Venezuelan Equine Encephalomyelitis Virus from a Vampire Bat Captured in Oaxaca, Mexico, 1970. Science 1972, 175 (4021), 546–547.

(75) Megger, D. A.; Philipp, J.; Le-Trilling, V. T. K.; Sitek, B.; Trilling, M. Deciphering of the Human Interferon-Regulated Proteome by Mass Spectrometry-Based Quantitative Analysis Reveals Extent and Dynamics of Protein Induction and Repression. Frontiers in Immunology. 2017. https://doi.org/10.3389/fimmu.2017.01139.

(76) Krapp, C.; Hotter, D.; Gawanbacht, A.; McLaren, P. J.; Kluge, S. F.; Stürzel, C. M.; Mack, K.; Reith, E.; Engelhart, S.; Ciuffi, A.; Hornung, V.; Sauter, D.; Telenti, A.; Kirchhoff, F. Guanylate Binding Protein (GBP) 5 Is an Interferon-Inducible Inhibitor of HIV-1 Infectivity. Cell Host Microbe 2016, 19 (4), 504–514.

(77) Braun, E.; Hotter, D.; Koepke, L.; Zech, F.; Groß, R.; Sparrer, K. M. J.; Müller, J. A.; Pfaller, C. K.; Heusinger, E.; Wombacher, R.; Sutter, K.; Dittmer, U.; Winkler, M.; Simmons, G.; Jakobsen, M. R.; Conzelmann, K.-K.; Pöhlmann, S.; Münch, J.; Fackler, O. T.; Kirchhoff, F.; Sauter, D. Guanylate-Binding Proteins 2 and 5 Exert Broad Antiviral Activity by Inhibiting Furin-Mediated Processing of Viral Envelope Proteins. Cell Reports. 2019, pp 2092–2104.e10. https://doi.org/10.1016/j.celrep.2019.04.063.

(78) Tretina, K.; Park, E.-S.; Maminska, A.; MacMicking, J. D. Interferon-Induced Guanylate-Binding Proteins: Guardians of Host Defense in Health and Disease. J. Exp. Med. 2019, 216 (3), 482–500.

(79) Yao, Z.; Jia, X.; Megger, D. A.; Chen, J.; Liu, Y.; Li, J.; Sitek, B.; Yuan, Z. Label-Free Proteomic Analysis of Exosomes Secreted from THP-1-Derived Macrophages Treated with IFN-α Identifies Antiviral Proteins Enriched in Exosomes. J. Proteome Res. 2019, 18 (3), 855–864.

(80) Ligtenberg, A. J. M.; Karlsson, N. G.; Veerman, E. C. I. Deleted in Malignant Brain Tumors-1 Protein (DMBT1): A Pattern Recognition Receptor with Multiple Binding Sites. Int. J. Mol. Sci. 2010, 11 (12), 5212–5233.

(81) Bikker, F. J.; Ligtenberg, A. J. M.; Nazmi, K.; Veerman, E. C. I.; van’t Hof, W.; Bolscher, J. G. M.; Poustka, A.; Nieuw Amerongen, A. V.; Mollenhauer, J. Identification of the Bacteria-Binding Peptide Domain on Salivary Agglutinin (gp-340/DMBT1), a Member of the Scavenger Receptor Cysteine-Rich Superfamily. J. Biol. Chem. 2002, 277 (35), 32109–32115.

(82) Hartshorn, K. L.; Ligtenberg, A.; White, M. R.; Van Eijk, M.; Hartshorn, M.; Pemberton, L.; Holmskov, U.; Crouch, E. Salivary Agglutinin and Lung Scavenger Receptor Cysteine-Rich Glycoprotein 340 Have Broad Anti-Influenza Activities and Interactions with Surfactant Protein D That Vary according to Donor Source and Sialylation. Biochem. J 2006, 393 (Pt 2), 545–553.

(83) McCarthy, M. K.; Weinberg, J. B. The Immunoproteasome and Viral Infection: A Complex Regulator of Inflammation. Front. Microbiol. 2015, 6, 21.

(84) Deshmukh, F. K.; Yaffe, D.; Olshina, M. A.; Ben-Nissan, G.; Sharon, M. The Contribution of the 20S Proteasome to Proteostasis. Biomolecules. 2019, p 190. https://doi.org/10.3390/biom9050190.

(85) Bayram, H. L.; Claydon, A. J.; Brownridge, P. J.; Hurst, J. L.; Mileham, A.; Stockley, P.; Beynon, R. J.; Hammond, D. E. Cross-Species Proteomics in Analysis of Mammalian Sperm Proteins. J. Proteomics 2016, 135, 38–50.

(86) Lai, R. C.; Tan, S. S.; Teh, B. J.; Sze, S. K.; Arslan, F.; de Kleijn, D. P.; Choo, A.; Lim, S. K. Proteolytic Potential of the MSC Exosome Proteome: Implications for an Exosome-Mediated Delivery of Therapeutic Proteasome. Int. J. Proteomics 2012, 2012, 971907.

(87) Neely, B. A.; Ferrante, J. A.; Mauro Chaves, J.; Soper, J. L.; Almeida, J. S.; Arthur, J. M.; Gulland, F. M. D.; Janech, M. G. Proteomic Analysis of Plasma from California Sea Lions (Zalophus Californianus) Reveals Apolipoprotein E as a Candidate Biomarker of Chronic Domoic Acid Toxicosis. PLOS ONE. 2015, p e0123295. https://doi.org/10.1371/journal.pone.0123295.

(88) Zhang, G.; Cowled, C.; Shi, Z.; Huang, Z.; Bishop-Lilly, K. A.; Fang, X.; Wynne, J. W.; Xiong, Z.; Baker, M. L.; Zhao, W.; Tachedjian, M.; Zhu, Y.; Zhou, P.; Jiang, X.; Ng, J.; Yang, L.; Wu, L.; Xiao, J.; Feng, Y.; Chen, Y.; Sun, X.; Zhang, Y.; Marsh, G. A.; Crameri, G.; Broder, C. C.; Frey, K. G.; Wang, L.-F.; Wang, J. Comparative Analysis of Bat Genomes Provides Insight into the Evolution of Flight and Immunity. Science 2013, 339 (6118), 456–460.

(89) Brunet-Rossinni, A. K. Reduced Free-Radical Production and Extreme Longevity in the Little Brown Bat (Myotis Lucifugus) versus Two Non-Flying Mammals. Mech. Ageing Dev. 2004, 125 (1), 11–20.

(90) Min-Oo, G.; Ayi, K.; Bongfen, S. E.; Tam, M.; Radovanovic, I.; Gauthier, S.; Santiago, H.; Rothfuchs, A. G.; Roffê, E.; Sher, A.; Mullick, A.; Fortin, A.; Stevenson, M. M.; Kain, K. C.; Gros, P. Cysteamine, the Natural Metabolite of Pantetheinase, Shows Specific Activity against Plasmodium. Exp. Parasitol. 2010, 125 (4), 315–324.

(91) Pitari, G.; Malergue, F.; Martin, F.; Philippe, J. M.; Massucci, M. T.; Chabret, C.; Maras, B.; Duprè, S.; Naquet, P.; Galland, F. Pantetheinase Activity of Membrane-Bound Vanin-1: Lack of Free Cysteamine in Tissues of Vanin-1 Deficient Mice. FEBS Lett. 2000, 483 (2-3), 149–154.

(92) Bartucci, R.; Salvati, A.; Olinga, P.; Boersma, Y. L. Vanin 1: Its Physiological Function and Role in Diseases. Int. J. Mol. Sci. 2019, 20 (16). https://doi.org/10.3390/ijms20163891.

(93) Tian, X.; Azpurua, J.; Hine, C.; Vaidya, A.; Myakishev-Rempel, M.; Ablaeva, J.; Mao, Z.; Nevo, E.; Gorbunova, V.; Seluanov, A. High-Molecular-Mass Hyaluronan Mediates the Cancer Resistance of the Naked Mole Rat. Nature 2013, 499 (7458), 346–349.

(94) Lin, C.-Y.; Kolliopoulos, C.; Huang, C.-H.; Tenhunen, J.; Heldin, C.-H.; Chen, Y.-H.; Heldin, P. High Levels of Serum Hyaluronan Is an Early Predictor of Dengue Warning Signs and Perturbs Vascular Integrity. EBioMedicine 2019, 48, 425–441.

(95) Alspach, E.; Lussier, D. M.; Schreiber, R. D. Interferon γ and Its Important Roles in Promoting and Inhibiting Spontaneous and Therapeutic Cancer Immunity. Cold Spring Harb. Perspect. Biol. 2019, 11 (3). https://doi.org/10.1101/cshperspect.a028480.

(96) Letko, M.; Miazgowicz, K.; McMinn, R.; Seifert, S. N.; Sola, I.; Enjuanes, L.; Carmody, A.; van Doremalen, N.; Munster, V. Adaptive Evolution of MERS-CoV to Species Variation in DPP4. Cell Rep. 2018, 24 (7), 1730–1737.

(97) Anthony, S. J.; Ojeda-Flores, R.; Rico-Chávez, O.; Navarrete-Macias, I.; Zambrana-Torrelio, C. M.; Rostal, M. K.; Epstein, J. H.; Tipps, T.; Liang, E.; Sanchez-Leon, M.; Sotomayor-Bonilla, J.; Aguirre, A. A.; Ávila-Flores, R.; Medellín, R. A.; Goldstein, T.; Suzán, G.; Daszak, P.; Lipkin, W. I. Coronaviruses in Bats from Mexico. J. Gen. Virol. 2013, 94 (Pt 5), 1028.

(98) Orsburn, B. C.; Jenkins, C.; Miller, S. M.; Neely, B. A.; Bumpus, N. N. In Silico Approach toward the Identification of Unique Peptides from Viral Protein Infection: Application to COVID-19. bioRxiv, 2020, 2020.03.08.980383. https://doi.org/10.1101/2020.03.08.980383.

(99) Bezstarosti, K.; Lamers, M. M.; Haagmans, B. L.; Demmers, J. A. A. Targeted Proteomics for the Detection of SARS-CoV-2 Proteins. bioRxiv, 2020, 2020.04.23.057810. https://doi.org/10.1101/2020.04.23.057810.

(100) Ihling, C.; Tänzler, D.; Hagemann, S.; Kehlen, A.; Hüttelmaier, S.; Arlt, C.; Sinz, A. Mass Spectrometric Identification of SARS-CoV-2 Proteins from Gargle Solution Samples of COVID-19 Patients. J. Proteome Res. 2020. https://doi.org/10.1021/acs.jproteome.0c00280.

(101) Gouveia, D.; Miotello, G.; Gallais, F.; Gaillard, J.-C.; Debroas, S.; Bellanger, L.; Lavigne, J.-P.; Sotto, A.; Grenga, L.; Pible, O.; Armengaud, J. Proteotyping SARS-CoV-2 Virus from Nasopharyngeal Swabs: A Proof-of-Concept Focused on a 3 Min Mass Spectrometry Window. J. Proteome Res. 2020. https://doi.org/10.1021/acs.jproteome.0c00535.

(102) Neely, B. A.; Carlin, K. P.; Arthur, J. M.; McFee, W. E.; Janech, M. G. Ratiometric Measurements of Adiponectin by Mass Spectrometry in Bottlenose Dolphins (Tursiops Truncatus) with Iron Overload Reveal an Association with Insulin Resistance and Glucagon. Front. Endocrinol. 2013, 4, 132.

(103) Shi, T.; Song, E.; Nie, S.; Rodland, K. D.; Liu, T.; Qian, W.-J.; Smith, R. D. Advances in Targeted Proteomics and Applications to Biomedical Research. Proteomics 2016, 16 (15–16), 2160–2182.

(104) Meyer, J. G.; Schilling, B. Clinical Applications of Quantitative Proteomics Using Targeted and Untargeted Data-Independent Acquisition Techniques. Expert Rev. Proteomics 2017, 14 (5), 419–429.

